# Early oligodendrocyte dysfunction signature in Alzheimer’s disease: Insights from DNA methylomics and transcriptomics

**DOI:** 10.1101/2025.08.28.672821

**Authors:** Katherine Fodder, Hannah M.G. Smith, Umran Yaman, Ignazio S. Piras, Megha Murthy, John Hardy, Tammaryn Lashley, Rohan de Silva, Dervis A. Salih, Conceição Bettencourt

## Abstract

Much research into the aetiology of Alzheimer’s disease (AD) has focused on neuronal cell types, while studies on the contribution of glial cells, particularly oligodendrocytes (OLGs), are only starting to emerge. Altered brain DNA methylation, an epigenetic modification that provides the interplay between genetics and environmental cues to tightly regulate gene expression, is well documented in AD. Yet, cell-type-specific investigations remain limited. Here, we examine the role of DNA methylation and OLGs in AD, and how such changes may impact gene expression. We performed weighted-gene correlation network analysis (WGCNA) on multiple brain omics AD datasets across species: human DNA methylation data from 4 brain regions, human brain single-nuclei RNA sequencing data and mouse brain RNA sequencing data. We compared AD-associated network modules enriched for OLG genes across AD brain regions, as well as with other neurodegenerative disease DNA methylation datasets. We identified a DNA methylation signature associated with AD, enriched for OLGs, and preserved across brain regions representing early and late AD pathology stages. Genes within this signature showed altered expression in AD OLGs, confirming cell-type specificity and relevance to AD. This OLG signature was also preserved in transgenic mice with early Aβ pathology and in other neurodegenerative diseases without Aβ pathology. We reveal a consistent pattern of OLG dysfunction spanning early to late stages of AD, across DNA methylation and gene expression. Our findings highlight OLG-associated DNA methylation changes as important in AD pathogenesis, and possibly in other neurodegenerative diseases, opening new avenues for therapeutic development.

## BACKGROUND

Alzheimer’s disease (AD) is the leading cause of dementia worldwide[1]. Although neurodegenerative diseases such as AD have been associated with grey matter and neuronal damage, there is evidence for decline and involvement of white matter and glial cells during disease progression[2–6]. Oligodendrocytes (OLGs), which are not well studied in neurodegenerative diseases, are glial cells in the CNS, responsible for the production, stability, and maintenance of myelin, the lipid-rich, multilamellar membrane which wraps around axons and enables fast transmission of electrical signals[7]. Early evidence of the disruption of myelin in AD suggests those regions of the cortex, such as the temporal and frontal lobes, that are myelinated later in development are more likely to present with AD pathology earlier[2,4,8]. This suggests that those regions that myelinate later are more vulnerable to pathogenic mechanisms which result in neurodegeneration. Moreover, brain imaging data has indicated that Aβ deposition may change white matter microstructure in early disease stages[9]. Although the precise role of OLGs in AD pathology is still unclear, there is evidence from human post-mortem studies showing alterations in the numbers and morphology of OLG lineage cells[3] as well as decreases in Olig2□+□cells[10], and an increased number of oligodendrocyte precursor cells (OPCs) in white matter lesions[11]. Morphological changes are also seen in OLGs derived from AD post-mortem brains, specifically decreased nuclear diameter in the parahippocampal white matter[12]. Genome-wide association studies (GWAS) have also implicated specific myelin/OLG-related genes in neurodegeneration, including the bridging integrator 1 (*BIN1*) gene, which is the second strongest genetic risk factor for late onset AD[13–15] and known to be largely expressed by mature OLGs in white matter tracts[16]. As well as observed changes in OLGs in AD, recent research has provided evidence OLG derived Aβ contributes to AD pathology, and that suppression of such production by OLGs in mice reduces AD associated pathology[17]. Transcriptomic analyses reveal gene expression changes in additional myelin-related/OLG genes in AD[18], further supporting the importance of myelination changes in AD. Interestingly, there are also genetic links between AD and some leukodystrophies, a rare group of inherited disorders characterised by loss of myelin[19]. For example, genetic variants in the AD associated gene *TREM2* are thought to be causative for an adult onset leukodystrophy[20,21]. Several other Mendelian leukodystrophy genes (e.g. *CSF1R* and *NOTCH3*) are also linked to AD[22]. This genetic overlap underscores the need to elucidate mechanisms of dysfunction of myelin in AD. Although OLG and myelin dysfunction in AD have historically been considered as consequences of neuronal pathology rather than causative factors, emerging evidence suggests that changes in myelin are an early feature of AD[3,23–26], indicating that oligodendrocyte dysfunction may play a contributory and early role in disease development.

DNA methylation, the most widely studied epigenetic modification, is crucial in the spatio-temporal control of gene expression, but has also been implicated in neurodegenerative disease[27–29]. In AD, epigenome-wide association studies (EWAS) utilising bulk brain tissue have identified multiple genes with DNA methylation changes associated with the disease and its pathological burden[30–33], and meta-analyses have identified significant changes across multiple brain regions[29,34–36]. More recently, various studies have investigated DNA methylation changes in glia and neurons separately, and have identified many AD-associated DNA methylation changes that had previously been detected in ‘bulk’ tissue but were indeed driven by changes in non-neuronal cells, including in OLGs[34,37]. However, detailed characterization of glial cell-type specific effects of dysregulated DNA methylation, particularly in OLGs, is limited.

In this study, we aimed to investigate dysregulation of OLG DNA methylation and their contribution towards AD. We utilised co-methylation network analysis, an agnostic systems biology approach to identify higher order relationships between biological entities, to uncover DNA methylation signatures derived from human post-mortem brain tissue that are associated with AD and relevant to OLG lineage cells. Following the identification of these AD-associated co-methylation OLG signatures, we investigated whether they were present in brain regions differently affected by AD pathology as well as in additional neurodegenerative diseases, such as primary tauopathies. Additionally, we integrated human brain single-nucleus RNA sequencing data and mouse brain gene co-expression networks to further investigate these OLG dysregulated signatures and whether they might be occurring early in the disease process. By bringing together multiple data modalities, we provide a multi-layered view of epigenetic and transcriptional changes supporting an important role for molecular dysregulation of OLGs early in the neurodegenerative process, an area which has so far been overlooked.

## METHODS

### 2.1 DNA methylation datasets and pre-processing

Data used to investigate DNA methylation signatures in AD was generated as described by Semick et al.[27]. This dataset comprised samples across multiple brain regions, including the cerebellum (CRB) (N = 67), which is affected last by AD pathology, while the hippocampus (HIPPO) (N = 65), entorhinal cortex (ERC) (N = 69), and dorsolateral prefrontal cortex (DLPFC) (N = 67) are all affected to varying degrees in AD, with the ERC and HIPPO among the earliest sites of pathology (i.e. representing advanced pathology) and the DLPFC becoming involved at later disease stages (i.e. representing earlier stages of the pathology compared with ERC and HIPPO) (**Table 1**). We also utilised the ROSMAP DNA methylation cohort (N = 530, DLPFC) in follow up analyses [30] (**Table 1)**

**Table 1.** Characterization of the AD datasets and selected regression models for input to downstream analyses.

| Dataset | AD cases and controls included after quality control | Mean age±SD (years) | Sex | Regression models used for covariate adjustment | Number (%) of methylation sites included for WGCNA | Soft-thresholding power used for WGCNA |
| --- | --- | --- | --- | --- | --- | --- |
| Primary multi-region Human DNA methylation discovery datasets |  |  |  |  |  |  |
| AD-HIPPO | AD (*N* = 16) | 81.8 ± 9.4 | 7M/9F | *~0+Sample_group+plate+DoubleN+sex+NeuNP+age+array* | 44,609 (20%) | 14 |
| | Controls (*N* = 47) | 61.7 ± 7.74 | 28M/19F | | | |
| AD-DLPFC | AD (*N* = 21) | 79.9 ± 9.46 | 11M/10F | *~0+Sample_group+NeuNP+age+sex+plate* | 53,826 (20%) | 14 |
| | Controls (*N* = 46) | 61.3 ± 7.56 | 27M/19F | | | |
| AD-ERC | AD (*N* = 19) | 79.3 ± 9.75 | 9M/10F | *~0+Sample_group+slide+DoubleN+Sox10P+age + 1/1 surrogate variables detected* | 52,296 (20%) | 12 |
| | Controls (*N* = 49) | 61.6 ± 7.61 ± 7.61 | 29M/20F | | | |
| AD-CRB | AD (*N* = 24) | 79.5 ±10.0 | 11M/13F | *~0+Sample_group+slide+sex+age+NeuNP+Double N* | 52,712 (20%) | 16 |
| | Controls (*N* = 43) | 60.6 ± 7.04 | 24M/19F | | | |
| Follow up Human DNA methylation and RNA sequencing datasets from overlapping brain donors |  |  |  |  |  |  |
| AD ROSMAP | AD (*N* = 201) | 85.4 ± 5.39 | 120M/81F | *DNA methylation model:*<br>*~0 + Sample_group + slide + bs_conversion + NeuN + DoubleN + array + PC1+ PC2*<br>*RNA sequencing model:* | NA | NA |
| | Controls (*N* = 329) | 87.3 ± 3.91 | 212M/117 F | | | |
|  |  |  |  | ~0 + disease + age + sex<br>+ pmi + sequencing batch |  |  |

AD, Alzheimer's disease; HIPPO, hippocampus; DLPFC, dorsolateral prefrontal cortex; ERC, entorhinal cortex; CRB, cerebellum; SD, standard deviation; F, females; M, males; NeuNP, estimated NeuN-positive nuclei proportion; DoubleN, estimated NeuN-/SOX10- nuclei proportion; Sox10P, estimated SOX10-positive nuclei proportion; WGCNA, weighted gene-correlation network analysis; PC1, 1st principal component; PC2, 2nd principal component; bs\_conversion, bisulfite conversion efficiency; pmi, post-mortem delay.

Beta-values ranging from 0 to 1 (approximately 0% to 100% methylated, respectively), were used to estimate the methylation levels of each CpG site using the ratio of intensities between methylated and unmethylated alleles. Data analysis was conducted using several R Bioconductor packages as previously described[38]. After importing raw data in the form of idat files, all datasets were subjected to pre-processing and quality control checks using minfi[39], wateRmelon[40], and ChAMP[41] packages. Probes were removed if they: (1) showed poor quality, (2) were cross reactive, (3) included common genetic variants, and (4) mapped to X or Y chromosome. Samples were also dropped if: (1) they presented with a high failure rate (≥□2% of probes), (2) the predicted sex did not match the phenotypic sex, and (3) they clustered inappropriately on multidimensional scaling analysis. Beta-values were normalised with ChAMP using the Beta-Mixture Quantile (BMIQ) normalisation method. M-values, computed as the logit transformation of beta-values, were used for all statistical analysis, as recommended by Du et al.[42], due to their reduced heteroscedasticity and improved statistical validity in comparison to beta-values for differential methylation analysis.

An overview of the methodological framework of this study is presented in **Fig. 1**. Following the approach described by Fodder et al.[43], and further detailed below, we conducted weighted-gene correlation network analysis (WGCNA). Prior to input into WGCNA, we carried out covariate adjustment using linear regression modelling to account for unwanted technical and biological effects (**Table 1**). Brain cell-type proportions were estimated using a cell-type deconvolution method and reference panel (NeuNP, NeuN-positive nuclei; DoubleN, NeuN−/SOX10− nuclei; Sox10P, SOX10-positive nuclei) described by Shireby et al[34]. Estimated cell-type proportions (NeuNP/DoubleN/Sox10P) were included as covariates whenever appropriate in region-specific regression models prior to network construction to mitigate confounding from cellular heterogeneity (**Table 1**).

**Fig. 1.**
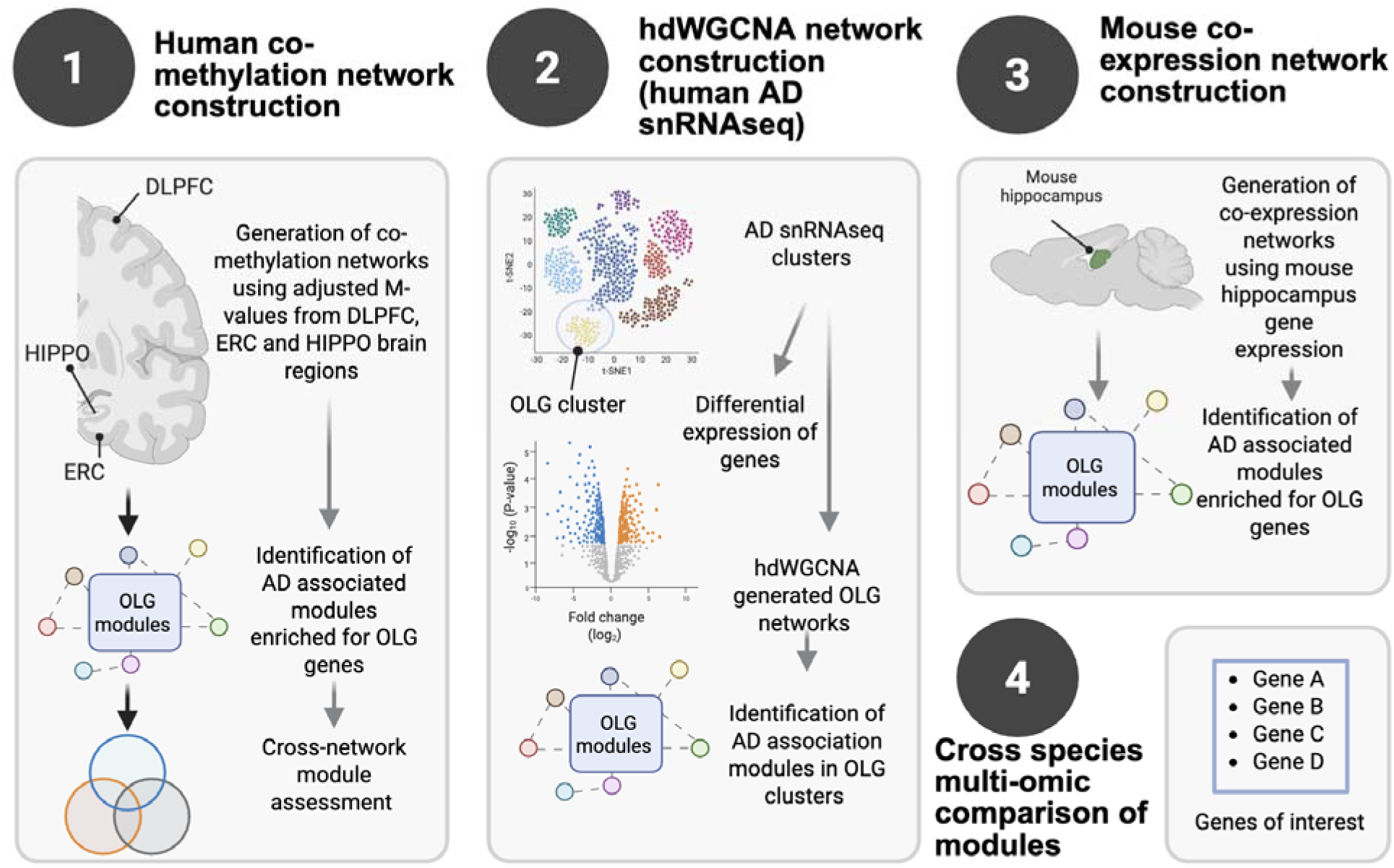
Overview of the methodological framework with a multi-omic network analysis approach to identify oligodendrocyte (OLG)-associated gene dysregulation in AD. 1) Human AD co-methylation networks were constructed using adjusted M-values from multiple human brain regions (e.g. DLPFC, ERC, and HIPPO. 2) Human AD single-nucleus RNA sequencing (snRNA-seq) data was then used to identify OLG clusters, where differentially expressed genes were identified. HdWGCNA was applied to build OLG-specific co-expression networks. 3) Mouse hippocampal gene expression data of AD models was then used to construct co-expression networks that were enriched for OLG signatures. 4) Modules identified from the three network types were compared to uncover genes present across multiple modules. DLPFC, dorsolateral prefrontal cortex; ERC, entorhinal cortex; HIPPO, hippocampus; AD, Alzheimer’s disease; OLG, oligodendrocyte; WGCNA, weighted gene-correlation network analysis; hdWGCNA, high-dimensional weighted gene-correlation analysis; snRNAseq, single-nucleus RNA sequencing.

### 2.2 Human brain gene expression

To investigate relationships between DNA methylation and gene expression, we analysed a bulk RNA-sequencing dataset (Table 1) comprising AD and control individuals from the ROSMAP cohort [30]. Raw count data were processed using the limma framework. Genes with low expression (maximum counts per million (CPM) < 1 across all samples) were excluded, normalization factors were calculated to account for differences in library size, and the voom function was applied to transform counts into log2-CPM values suitable for linear modelling. Linear models were fitted to identify associations with disease statusand adjust for relevant covariates (Table 1), based on singular value decomposition (SVD) analysis.

To investigate relationships between DNA methylation and gene expression, relevant modules identified in the discovery dataset were evaluated in an independent cohort for which matched DNA methylation and bulk RNA-sequencing data were available. For each module of interest, CpG sites assigned to that discovery module were identified in the independent DNA methylation dataset, thereby projecting the modules into the follow-up cohort. Module eigengenes, i.e. first principal component, were then calculated for each sample using the methylation levels of these CpG sites, generating a single summary measure representing the overall methylation pattern of the module. Expression levels of genes to which CpGs in each module mapped were extracted from the matched RNA-seq dataset. Pearson correlations were then computed between methylation-derived module eigengenes and gene expression levelsto assess the relationship between coordinated methylation and transcriptional variation.

We used human brain single-nucleus RNA sequencing (snRNAseq) data from the previously published and publicly available ROSMAP study[44]. We used filtered cell counts from 24 individuals showing little or no pathology and 24 showing mild to severe AD pathology (*N* = 48 total). Pathology stage was also assigned to each individual as ‘no-pathology’, ‘early-pathology’ or ‘late-pathology’ as described by Mathys et al.[44]. Filtered data consisted of 70,634 droplet-based single-nucleus covering a total of 17,926 genes. We used Seurat[45] to normalise the data with the function “NormalizeData” with the option “LogNormalize”, using a scale factor of 10,000. We then assigned cell cluster identities as described by Mathys et al.[44], and considered OLG subclusters (Oli0,1,3,4 and 5) only.

### 2.3 Mouse brain gene expression

To investigate early disease changes, we also examined mouse brain gene expression to complement the human post-mortem end stage disease findings. Such mouse models represent the early-stage of amyloid pathology but no tau pathology. The gene expression dataset was generated using AD mouse models as described by Salih *et al*.[46], Matarin *et al.*[47]. In brief, four transgenic mouse lines which develop Aβ pathology at varying rates were included: *APP, PSEN1,* homozygous *APP/PSEN1* and heterozygous *APP/PSEN1* alongside age-matched wild-type mice (**Supplementary Table S1**). For each transgenic line, at least three mice were used, except for *APP* mice at 4 months, where only two mice were available (**Supplementary Table S1**). A minimum of seven littermate controls (*WT*) from the original parental lines were also included. The hippocampi of male mice at the age of 2, 4, 8 and 18 months were used for transcriptomic analysis. Procedures involving mice were conducted in accordance with local ethical agreement, and in compliance with the UK Animals Act (1986).

As described by Salih *et al.*[46], RNA-seq library preparation and sequencing were performed by Eurofins Genomics (Ebersberg, Germany). Briefly, RNA strand-specific libraries were created using TruSeq Stranded mRNA Library Preparation Kit, Illumina according to the manufacturer’s instructions. Raw sequencing data were processed using RTA version 1.18.64, and FASTQ files were generated using bcl2fastq-1.8.4. Low-quality base pairs and adaptors were removed using Trim Galore (Babraham Bioinformatics). Transcripts were quantified using Salmon[48] and gene level quantification, expressed as transcripts per million (TPM), was obtained using the tximport package in R[49]. Gene expression values were transformed by applying log2 (normalised counts+1), and lowly expressed genes, i.e. those with an average detection level of below 1.5 TPM were excluded, reducing the dataset from 48,525 to 15,453 genes.

### 2.4 Weighted gene correlation network analysis (WGCNA)

We used a systems biology approach to identify co-methylation and co-expression modules, i.e. clusters of highly correlated DNA methylation sites (CpGs) or genes. For this we have used weighted gene-correlation network analysis (WGCNA[50]) for the four human bulk brain tissue DNA methylation datasets (**Table 1**) and for the mouse gene expression dataset (**Supplementary Table S1**). For all DNA methylation datasets, we excluded those CpGs not mapping to genes and selected only those showing the highest variance (top 20%) across individuals within each dataset (**Supplementary Tables 2.1-2.3**). The percentage of total methylation sites and number of methylation sites included for each analysis is given in **Table 1**, as is the soft-thresholding power used for network construction. Selected CpGs are representative of feature distribution in the original CpGs background pool (**Supplementary Table S3**). We used covariate adjusted M-values for the selected CpGs as input for this analysis. For the mouse expression data, following sample clustering, three outliers were removed leaving 83 samples for the network analysis. A soft-thresholding power of 6 was used. To construct all networks, pairwise DNA methylation/expression correlations were calculated to create a similarity matrix, as previously described[50].

Module membership (MM) was then reassigned for each network using the applyKMeans function of the CoExpNets package[51]. CpGs/genes with high module membership (MM) are considered network hubs, reflecting high intramodular connectivity[50]. This measure indicates centrality within the network structure. Gene significance was also considered, which here represents the correlation between the methylation levels at a specific site and the disease status[50].

After co-methylation/expression modules were identified, we carried out module-trait correlation analysis (as part of the WGCNA workflow[50]) to understand the relationship between co-methylation/expression modules and disease traits. For the human co-methylation modules, this association was carried out with AD status (i.e. AD or control sample). Whilst for the mouse co-expression modules association was based on a semi-quantitative score based on Aβ pathology severity, ranging from 0-5 obtained as previously described[46]. We applied Bonferroni multiple testing correction, i.e. corrected correlation p-value over the number of modules detected, to minimize the finding of false-positives.

### 2.5 Cell-type enrichment analyses for co-regulation modules

Cell-type enrichment analysis for the human co-methylation and mouse co-expression modules was conducted using the EWCE package and the associated single-cell mouse transcriptomic dataset[52,53]. This analysis aimed to determine whether the genes within each co-methylation/co-expression module were enriched for markers of a specific cell type. Overrepresented cell types were identified using p-values derived from 1000 iterations. For the human co-methylation modules we additionally checked this enrichment using an OLG gene list from human snRNA-sequencing data generated by Mathys et al.[44] and curated by Piras et al.[54]. We used Fisher’s exact test to confirm whether OLG modules identified through the EWCE package were also statistically significantly enriched for genes within this OLG list. For both EWCE and Fisher’s exact tests, enrichment was assessed relative to the background set of all genes represented in the corresponding network input (i.e. all genes mapped by CpGs retained after selection for WGCNA).

### 2.6 High dimensional WGCNA (hdWGCNA)

To confirm the cell-type specificity and the OLG relevance of the identified co-methylation signatures, we also utilised hdWGCNA to identify co-expression networks within the high-dimensional snRNA-seq dataset. Using cells belonging to the five OLG clusters (Oli_0-5 described above), we restricted our analysis to genes that were expressed in at least 1% of cells (total genes = 9,254). Following the pipeline described by Morabito et al.[55], we generated metacells, which aggregate similar cells within each group (OLG subclusters) and averaged their expression profiles, thereby reducing noise and improving module detection. The resulting metacell expression matrix was then log-normalized, before soft-thresholding was carried out to identify a power of 8 for network construction.

### 2.7 Preservation Analysis

To identify which co-methylation modules in each of AD networks were preserved (i.e. shared) or perturbed (i.e. unique) in the other brain regions, and against additional AD and other neurodegenerative disease datasets, we employed module preservation analysis, as described by Langfelder et al.[56]. For each network (taken as the “reference dataset”), module preservation in the other datasets (the “test data”) was calculated using the modulePreservation function implemented in WGCNA, carried out with 200 permutations for each analysis. As a measurement of module preservation, we used the Z-summary statistic (a composite measure to summarise multiple preservation statistics). A Z-summary greater than 10 indicates a strong preservation of a module in the “test data”, a Z-summary of between 2 and 10 indicates moderate preservation, and a Z-summary less than 2 indicates no preservation. To investigate preservation, we used five additional complementary DNA methylation datasets derived from AD, frontotemporal lobar degeneration (FTLD) and movement disorders (MD), which are summarised in **Table 2**.

**Table 2.** Pathological and demographic characteristics of DNA methylation datasets used for preservation analysis.

| Dataset | Disease subtypes and controls (N after quality control) | Mean age±SD (years) | Sex | Regression models used for covariate adjustment |
| --- | --- | --- | --- | --- |
| FTLD1<br>(Frontal Cortex)<br>Fodder et al.[43] | FTLD-TDP (N=15) | 70.07±5.59 | 7M/8F | $\sim 0 + \text{disease} + \text{age} + \text{sex} + S_{OX10P} \text{ proportions} + \text{DoubleN proportions} + \text{array}$ (0 surrogate variables detected) |
| | Controls (N=8) | 75.75±5.63 | 3M/5F | |
| FTLD2<br>(Frontal Cortex)<br>Menden et al.[57] | FTLD-TDP/FTLD-tau (N=34) | 63.18±7.92 | 14M/20F | $\sim 0 + \text{disease} + \text{age} + \text{sex} + S_{OX10P} \text{ proportions} + \text{DoubleN proportions} + \text{array} + \text{slide}$ (0 surrogate variables detected) |
| | Controls (N=14) | 78.43±11.76 | 5M/9F | |
| FTLD3<br>(Frontal Cortex)<br>Weber et al.[58] | FTLD-Tau (PSP) (N=93) | 71.16±5.32 | 54M/39F | $\sim 0 + \text{disease} + \text{age} + \text{sex} + S_{OX10P} \text{ proportions} + \text{DoubleN proportions} + \text{array} + \text{slide} + \text{surrogate variable}$ (1/1 surrogate variables detected) |
| | Controls (N=70) | 76.17±7.93 | 45M/25F | |
| AD<br>(Dorsolateral Prefrontal Cortex)<br>De Jager et al.[30] | AD (N=201) | 85.4 ± 5.39 | 120M/81F | $\sim 0 + \text{disease} + \text{slide} + \text{bs\_conversion} + \text{NeuNP proportion} + \text{DoubleN proportion} + \text{array} + \text{PC1} + \text{PC2}$ |
| | Controls (N=329) | 87.3 ± 3.91 | 212M/117F | |
| MD<br>(Frontal lobe white matter)<br>Murthy et al.[59] | Controls (N=17) | 72.35 ± 4.47 | 8M/9F | $\sim 0 + \text{disease} + \text{age} + \text{sex} + \text{PMI} + \text{NeuNP proportion} + \text{DoubleN proportion} + \text{slide} + \text{array}$ |
| | MSA (N=17) | 67.71 ± 6.47 | 9M/8M | |
| | PD (N=17) | 68.06 ± 3.67 | 8M/9F | |
| | PSP (N=17) | 65.35 ± 3.75 | 9M/8F | |

### 2.8 Genetic enrichment assessment

To investigate whether any of the co-methylation modules we produced during the analysis described above were enriched for genes known to confer genetic risk of AD, we utilised MAGMA-Generalized Gene-Set Analysis of GWAS Data[60]. MAGMA uses a multiple regression framework to enable powerful and efficient gene-and gene-set level analysis.

## RESULTS

### 3.1 Human brain co-methylation network analysis reveals significant oligodendrocyte dysregulation associated with AD

We generated co-methylation networks for four brain regions affected by different degrees of AD pathology (**Supplementary Fig. S1**), with the HIPPO representing the brain region displaying pathology earlier in AD, followed by the ERC and DLPFC, and the CRB, which is usually spared from typical AD pathology. For the DLPFC, ERC, and HIPPO and CRB networks, 6/16 (*p*□<□0.003, 0.05/16 modules), 17/25 (*p*□<□0.002, 0.05/25 modules), 7/18 (*p*□<□0.003, 0.05/18 modules) and 4/6 (*p*□<□0.008, 0.05/6 modules) co-methylation modules, respectively, were found to be associated with the disease status (i.e. AD vs control) (**Supplementary Fig. S1, Supplementary Table S3.1-3.3**). We next analysed which of these modules were enriched for OLG/OPC relevant genes, and identified one module in each of the DLPFC, ERC and HIPPO networks, the DLPFC-greenyellow, ERC-tan and the HIPPO-grey60 (**Fig. 2, Supplementary Figures S1-S2, Supplementary Tables S4.1-4.3**). For the CRB networks, we did not find significant enrichment for OLGs/OPCs (**Supplementary Fig. S1B**). Despite not the focus of this work, as expected, EWCE additionally identified modules enriched for neuronal (e.g. DLPFC-green) and microglial (e.g. DLPFC-salmon) genes across networks (**Supplementary Figure S1**), several of which were also associated with AD status, consistent with the previously established involvement of these cell types in AD pathology. All OLG/OPC enriched modules (DLPFC-greenyellow, ERC-tan and the HIPPO-grey60) also showed significant positive correlation between module membership and gene-level significance for AD status (r = 0.69, r = 0.91, r = 0.53; p < 1×10^−138^; respectively), and a positive correlation with disease status, i.e. increased DNA methylation levels in AD compared to controls (**Supplementary Fig. S1**).

**Fig. 2.**
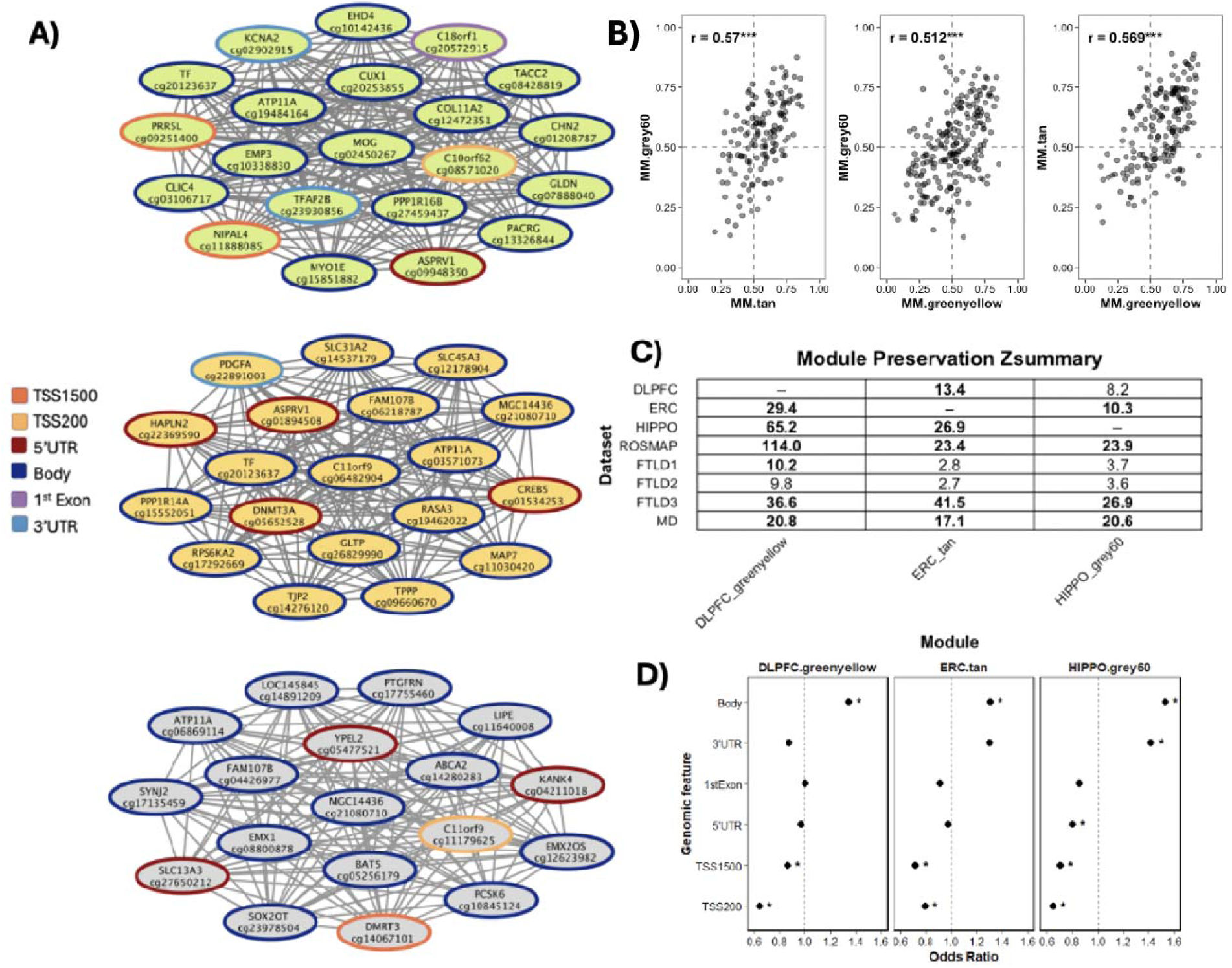
Human brain co-methylation network analysis shows significant dysregulation of OLG signatures across brain regions in AD. A) Top 25 genes with highest module membership across the DLPFC-greenyellow, ERC-tan and HIPPO-grey60 modules (top to bottom). Edge surrounding the genes indicates the feature of the CpG mapping to each gene as given by the legend. B) Scatterplots show pairwise relationships between CpG-level MM values from the grey60, tan, and greenyellow modules with Pearson correlation coefficients (r) and significance indicated (*** p < 0.001). C) Module preservation scores between co-methylation modules (DLPFC, ERC and HIPPO), ROSMAP (AD), FTLD, and MD test data. Bold values indicate high preservation (Zsummary > 10); regular text indicates moderate preservation (2 < Zsummary ≤ 10). D) Enrichment of methylation site features within 3 AD-associated modules of interest. Asterisks indicate significant enrichment (p < 0.05). AD, Alzheimer’s Disease; FTLD, Frontotemporal lobar degeneration; HIPPO, hippocampus; DLPFC, dorsolateral prefrontal cortex; ERC, entorhinal cortex; MD, movement disorder; MM, module membership; TSS200, transcription start site −200; TSS1500, transcription start site −1500-200; 3’UTR, 3 prime untranslated region; 5’UTR, 5 prime untranslated region.

Hub gene analysis, which identifies in this case the most strongly connected CpGs and related genes for each OLG-enriched module, reinforced the relevance for OLG lineage cells in AD-associated modules. The top hub gene (highest module membership (MM)) for the DLPFC-greenyellow module was *MOG* (myelin oligodendrocyte glycoprotein) (**Fig. 2A, Supplementary Table S4.1)**, a well described marker of mature OLGs[61], which has also been found differentially methylated in chronic demyelinated lesions in multiple sclerosis[62]. The CpG with the second highest MM mapped to *ATP11A* (ATPase phospholipid transporting 11A), a gene which is more highly expressed in OLGs compared to other brain cell types (<u>ATP11A Tissue Expression</u>) (**Supplementary Table S4.1),** and is mutated in hypomyelinating leukodystrophy[63]. We noted that 26 CpGs within this greenyellow module mapped to this gene *ATP11A* (**Supplementary Table S4.1**).

The top hub gene for the ERC-Tan module, with the two CpGs with highest MM, was *C11ORF9* (also known as *MYRF*) (**Fig. 2A, Supplementary Table S4.2**), which codes for myelin regulatory factor, an important driver of post-OLG differentiation myelination and myelin maintenance[64]. In a previous study using gene expression and proteomic networks, *MYRF* was found to be a key driver of AD-associated disrupted OLG signatures[65]. The *ATP11A* gene, which was also highlighted in the DLPFC-greenyellow module, has 12 CpGs within this tan module (**Supplementary Table S4.2**). A CpG mapping to *MBP* (a well described marker of mature OLGs) also presents with high MM within the module (**Supplementary Table S4.2**).

The hub gene for the HIPPO-grey60 module was *MGC14436,* according to the Illumina DNA methylation array manifest, which maps to chromosome 12 and is not well characterised. However, further inspection of the genomic location of this CpG revealed that it lies within the region of the *FAM222A* gene, which encodes a protein previously reported to build up in Aβ plaques in AD[66], and has been shown to contain genetic variants associated with AD risk[26]. The gene has also been reported to be co-expressed with several other AD related genes[67]. This disease-associated OLG module also contained 27 CpGs mapping to the gene *ATP11A* (**Supplementary Table S4.3)**, as we had seen with the ERC-tan and the DLPFC-greenyellow module. Full details of all CpGs within each module and corresponding MM information are given in **Tables S4.1-4.3.**

We also examined similarities across brain regions for the OLG-enriched AD co-methylation modules. Pairwise comparisons of module membership (MM) values revealed moderate and highly significant positive correlations between regions (**Fig. 2B**). We also utilised module preservation statistics to evaluate how well the co-methylation modules identified in one region were preserved in the others. The DLPFC-greenyellow module showed strong preservation in both ERC and HIPPO datasets (Z.summary > 10), as did the ERC-tan module in the DLPFC and the HIPPO networks. The HIPPO-grey60 module was preserved in ERC and DLPFC, albeit only moderately in DLPFC (**Fig. 2C**). Looking in more detail at which CpGs were present across all three, we found that 74 CpGs belonged to all DLPFC-greenyellow, ERC-tan and HIPPO-grey60 modules, mapping to 54 distinct genes (**Supplementary Table S5**). Notably, 8/74 probes mapped to the gene *ATP11A* (highlighted above). Other genes containing multiple probes within this set included *INPP5A* (to which 5/74 CpGs mapped), *RASA3* (3/74) and *FAM107B* (3/74).

To further assess the reproducibility of the co-methylation modules, we evaluated their preservation in an independent bulk AD DNA methylation dataset from the ROSMAP study (derived from prefrontal cortex tissue). All three modules showed strong preservation, suggesting that they represent robust biological signals (**Fig. 2C**). We also investigated whether this preservation extended to other neurodegenerative diseases, namely several frontotemporal lobar degeneration (FTLD) subtypes we previously investigated ranging between FTLD-tau and FTLD-TDP pathology[43] and a DNA methylation dataset derived from the white matter (highly enriched for OLGs) of multiple system atrophy (MSA), progressive supranuclear palsy (PSP) and Parkinson’s disease (PD) samples[59]. Both the DLPFC-greenyellow and the ERC-tan AD-associated modules (OLG enriched), were highly preserved (the most preserved of all co-methylation modules) in the FTLD3 dataset (composed of sporadic primary tauopathy cases - PSP) and moderately preserved in the FTLD1 and FTLD2 dataset (mostly TDP-43 proteinopathies) (**Fig. 2C**). All three modules were also highly preserved in the white matter MSA/PSP/PD dataset (**Fig. 2C**), indicating that in addition to AD these dysregulated OLG signatures are also highly relevant to other neurodegenerative diseases with multiple underlying pathologies.

We also checked to see if there was any enrichment of a specific CpG feature (i.e. where within the gene the CpG was located) within these modules. All three modules showed significant enrichment of CpGs located within gene bodies compared with CpGs assigned to other modules in the co-methylation network (**Fig. 2D**). Although the role of DNA methylation within the gene body is less well characterised compared to that in the promoter regions (where increased DNA methylation is often associated with a decrease in gene expression), our data points towards the importance of gene body DNA methylation dysregulation in AD.

We checked whether any of the modules were linked to AD risk genes using MAGMA and associated GWAS summary statistics[14], and found no enrichment within the three modules (data not shown), suggesting that known AD-associated genetic effects are unlikely to be driving the OLG dysregulation we report above.

### 3.2 AD-associated oligodendrocyte co-methylation modules show coordinated transcriptional regulation

To further explore the relationship between DNA methylation changes within these modules and transcriptional regulation, we analysed an independent cohort with matched DLPFC bulk DNA methylation and RNA-seq data. We projected the DLPFC-greenyellow co-methylation module into this follow-up methylation dataset **(Fig. 3A**). The projected module eigengene showed a significant association with Braak stage (**Fig. 3B**), indicating that coordinated methylation variation within this module tracks increasing AD pathology in bulk brain tissue.

**Fig. 3.**
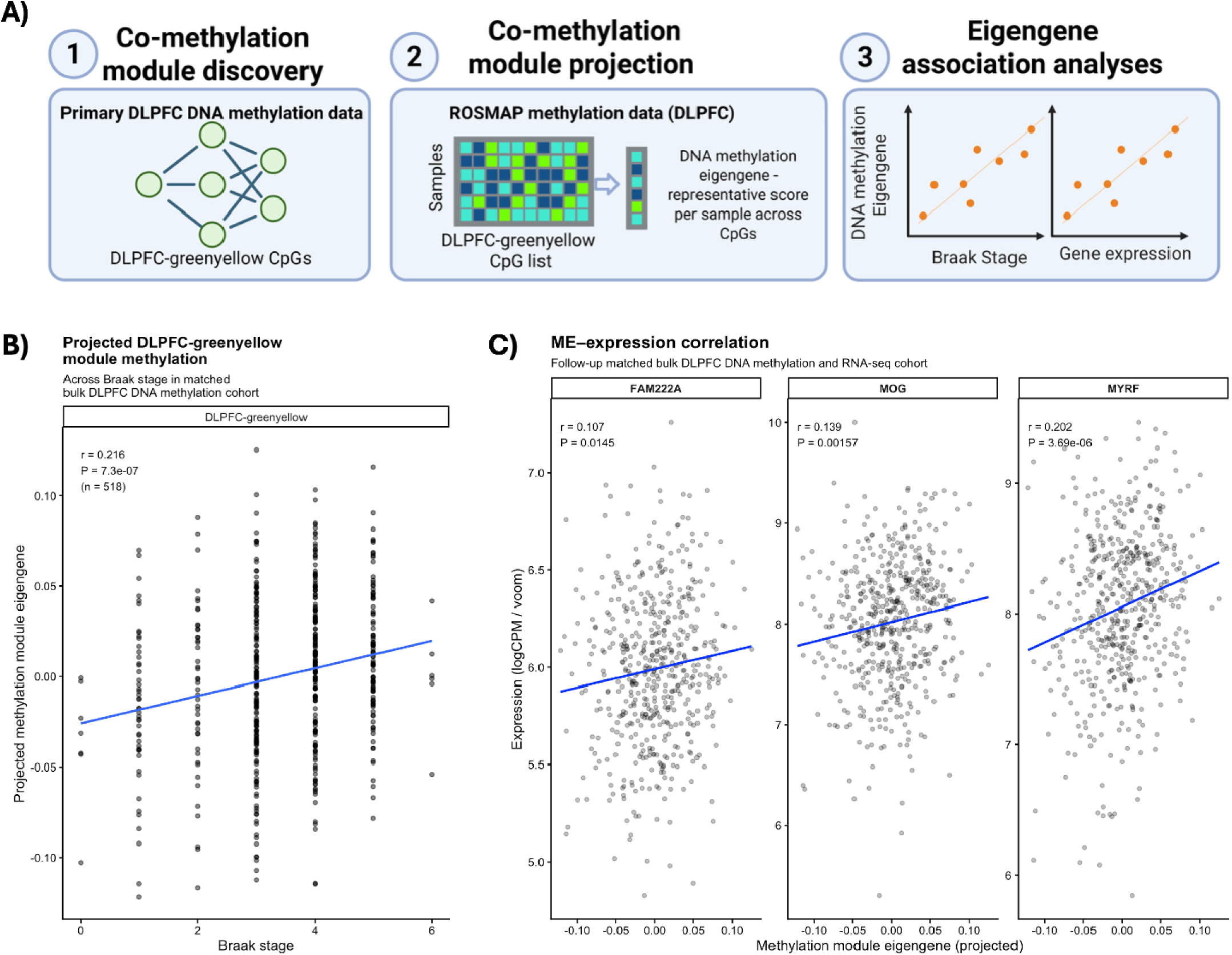
Projected co-methylation module genes show coordinated methylation–expression coupling and are associated with AD pathology. A) Overview of the analytical workflow. Co-methylation modules were first identified from DNA methylation data in the discovery dorsolateral prefrontal cortex (DLPFC) cohort using network analysis. Module CpGs were then projected onto an independent DLPFC DNA methylation dataset from the ROSMAP cohort to calculate a representative **module eigengene** (1st principal component) for each sample. Finally, associations between the projected module eigengene and neuropathological or transcriptional measures were evaluated. B) Association between the projected DLPFC-greenyellow module eigengene and Braak stage in an independent bulk DLPFC DNA methylation cohort. The module was defined in the multi-region discovery dataset and projected into the follow-up cohort using the same CpG set to calculate module eigengene values. Pearson correlation statistics are shown. C) DNA methylation–expression correlations in the matched bulk DLPFC cohort. Scatter plots show correlations between methylation-derived module eigengene values and expression of hub genes from the co-methylation modules (*FAM222A, MOG, MYRF*) in RNA-seq data from the same donors. Lines represent linear regression fits; Pearson r and p-values are indicated. DLPFC, dorsolateral prefrontal cortex; ME, module eigengene.

We next examined whether DNA methylation variation captured by this module was associated with transcriptional changes. This analysis revealed strong transcriptional correspondence for the projected DLPFC-greenyellow module, with over 50% of module genes showing significant correlations between gene expression and the module eigengene (FDR < 0.05) (**Supplementary Table S6**). Expression levels of hub genes from the co-methylation modules (*FAM222A*, *MOG*, and *MYRF*) also showed significant positive correlations with the module eigengene (**Fig. 3C)**. Together, these findings indicate that coordinated methylation variation within this module is reflected in the transcriptional output of key oligodendrocyte genes with a positive association between DNA methylation and expression levels.

### 3.3 Oligodendrocyte co-methylation module genes show mostly upregulation in AD OLGs

To assess whether genes within the AD-associated OLG co-methylation modules show cell-type–specific transcriptional changes during disease progression (early and late pathology), we examined their differential expression patterns in a human OLG single-nucleus RNA-sequencing (snRNA-seq) dataset (**Fig. 4A**). Genes from these modules were differentially expressed across several OLG subclusters, with the greatest number of expression changes observed in the Oli3 and Oli0 clusters. Notably, Mathys et al. previously described the Oli0 subcluster as being enriched for cells derived from samples with high amyloid burden and cognitive decline [44].

**Fig. 4.**
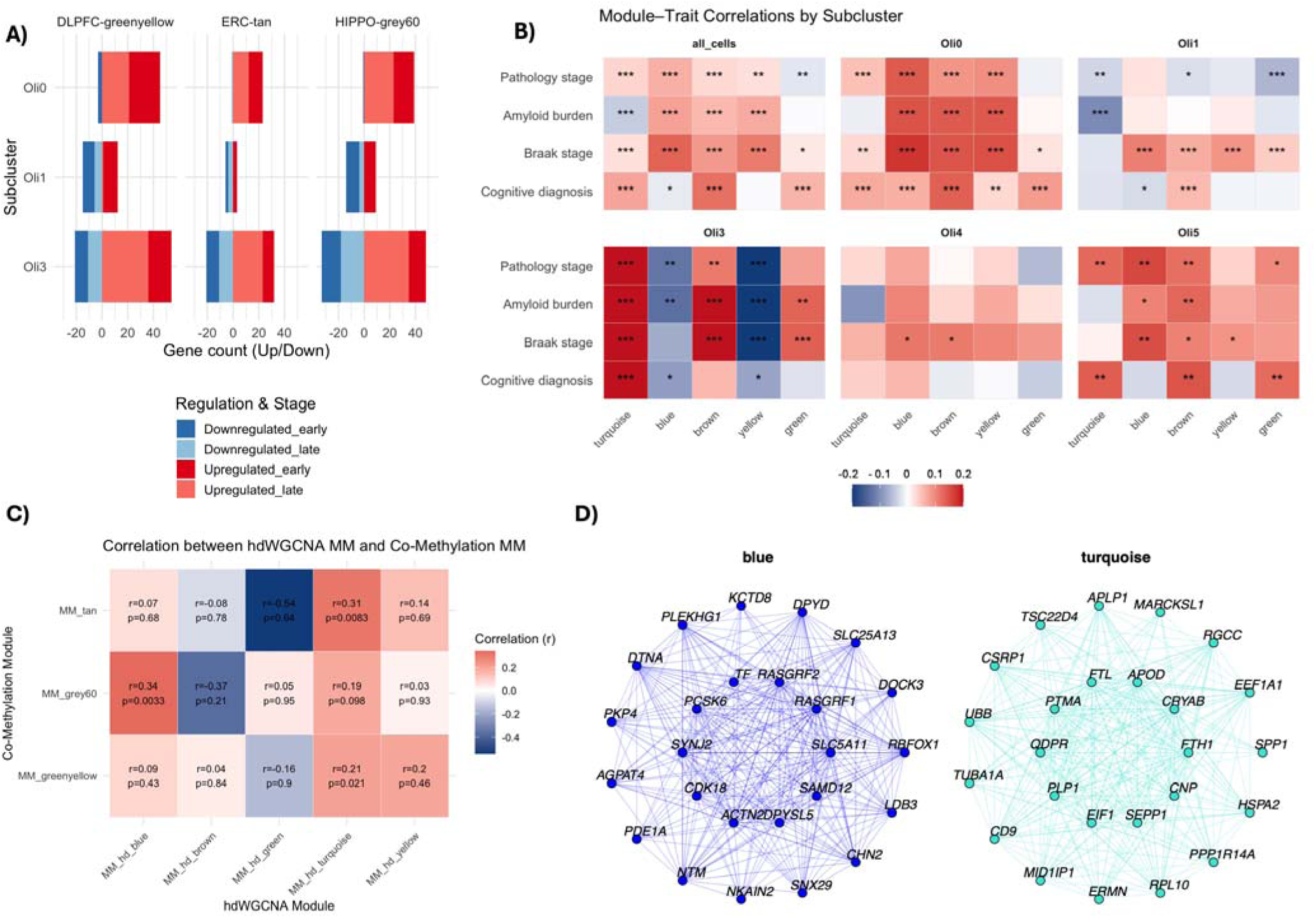
Co-expression network analysis of human oligodendrocyte snRNAseq clusters. A) Differential expression of genes belonging to DLPFC-greenyellow, ERC-tan, and HIPPO-grey60 co-methylation modules across oligodendrocyte (OLG) subclusters in snRNA-seq data. Bars represent the number of upregulated (red) and downregulated (blue) genes in early and late pathology comparisons (as defined by Mathys et al[44].) B) Heatmap showing correlations between module eigengenes of hdWGCNA co-expression modules and neuropathological traits (pathology stage, amyloid burden, Braak stage, and cognitive diagnosis (a clinical consensus on cognitive status at time of death ranging between 1 (no cognitive impairment) and 6 (dementia)) for all oligodendrocyte clusters and for individual subclusters (Oli0–Oli5). Red indicates positive and blue negative correlations. Significance is indicated: *p* < 0.05 (\**), p < 0.01 (**), p < 0.001 (\*\*\**). C) Heatmap of Pearson correlations between module membership (MM) values of three disease-associated co-methylation modules (ERC-tan, HIPPO-grey60 and DLPFC-greenyellow) and MM in each hdWGCNA module. D) Network plots of two hdWGCNA modules (blue and turquoise), showing connections amongst top 15 hub genes. hdWGCNA, high-dimensional weighted gene correlation network analysis; MM, module membership; DLPFC, dorsolateral prefrontal cortex; ERC, Entorhinal cortex; HIPPO, hippocampus; snRNAseq, single nucleus RNA sequencing.

Across subclusters, a large proportion of module genes showed increased expression across disease stages (i.e. both early and late). This predominance of upregulated genes is in line with findings from bulk tissue DNA methylation-gene expression patterns and with expected direction of effect liked to increased methylation in CpGs within gene body regions [67], which we found across all three OLG AD co-methylation modules. Among the genes most strongly upregulated within the Oli3 cluster were several present within our co-methylation modules, including *QDPR* and *CRYAB* (**Supplementary Fig. S3**), both of which have also been reported by Mathys et al. to show increased protein abundance in white matter from individuals with AD [44].

We also constructed gene co-expression modules from the oligodendrocyte snRNA-seq dataset. Using hdWGCNA, we identified 5 modules within the OLG clusters: hd-green, hd-brown, hd-turquoise, hd-blue and hd-yellow. All modules showed significant association with at least one pathological trait (i.e. amyloid, Braak stage and/or cognitive decline) within one subcluster of OLGs (**Fig. 4B**). Module–trait correlations were prominent within the Oli3 and Oli0 subclusters, where hdWGCNA module eigengenes showed strong associations with amyloid burden, Braak stage, and cognitive diagnosis (**Fig. 4B**). As we did between co-methylation modules across brain regions, we looked at correlation between module membership within co-methylation OLG modules and snRNAseq gene expression modules. The snRNAseq hd-turquoise module showed significant positive correlations with the DLPFC-greenyellow and ERC-tan modules (**Fig. 4B**), and non-significant but positive correlations with the HIPPO-grey60 module. The snRNAseq hd-blue module was significantly correlated with the HIPPO-grey60 module. We next looked at which genes showed high MM across co-methylation and co-expression modules (**Supplementary Table S7.1-7.2**). The gene with the second highest MM in the hd-turquoise module was *CRYAB,* which was also part of the DLPFC-greenyellow module (**Supplementary Table S4.1**). Other genes with high MM in the hd-turquoise module (top 5%) included *QDPR* (**Fig. 4C**), which was included in all three co-methylation modules (**Supplementary Tables S4.1-4.3**).

### 3.4 AD-associated OLG co-methylation networks show dysregulation of gene expression in mouse models of early AD stages

To complement human post-mortem brain tissue derived datasets and be able to infer the relevance of dysregulated OLG signatures at an early stage of AD pathology, we have used gene co-expression networks using a dataset derived from early-stage AD mouse models. A total of 10 mouse brain co-expression modules were identified and then related to sample traits: Aβ pathology, age and genotype (**Supplementary Fig. S4A**). Three modules were significantly positively associated with Aβ pathology (p ≤ 0.005 (0.05/10)), indicating that the expression of genes within these modules significantly increases with increasing pathology. We identified the pink mouse co-expression module as significantly associated with AD-related pathology and enriched for OLGs (**Supplementary Fig. S4B**).

Consistent with OLG involvement, well-known OLG marker genes were present across the mouse pink co-expression module, including *Mobp, Plp1, Fa2h, Sox10* and *Mag*, and hub genes included *Myo1d, Clic4, Ttyh2, Pacs2, Arhgef10, Tmem88b, Carns1, Cntn2,* and *Scd1* (**Fig. 5A, Supplementary Table S8**). There was a significant overlap of the mouse pink module genes with all three human OLG-relevant AD co-methylation modules (**Supplementary Fig. S4C**), and the hub genes of the ERC-tan and DLPFC-greenyellow modules, *Mog* and *Myrf* were also present within the pink module. The gene *Fam222A* (possible HIPPO-grey60 hub gene) was also present in this module. Furthermore, many of the overlapping genes present with high MM in both human and mice networks (**Fig. 5B**), suggesting convergence on shared OLG-related gene networks across species and molecular layers. Additionally, because these mice do not have tau pathology, this data suggests that dysregulation of these OLG signatures could occur at an early stage of AD pathology before tau pathology develops. Interestingly, because these OLG signatures are highly preserved in primary tauopathies and alpha-synucleinopathies (**Fig 2.C**), this suggests that their dysregulation may also be independent of amyloid pathology. A summary outline of these findings is shown in **Fig. 6**.

**Fig. 5.**
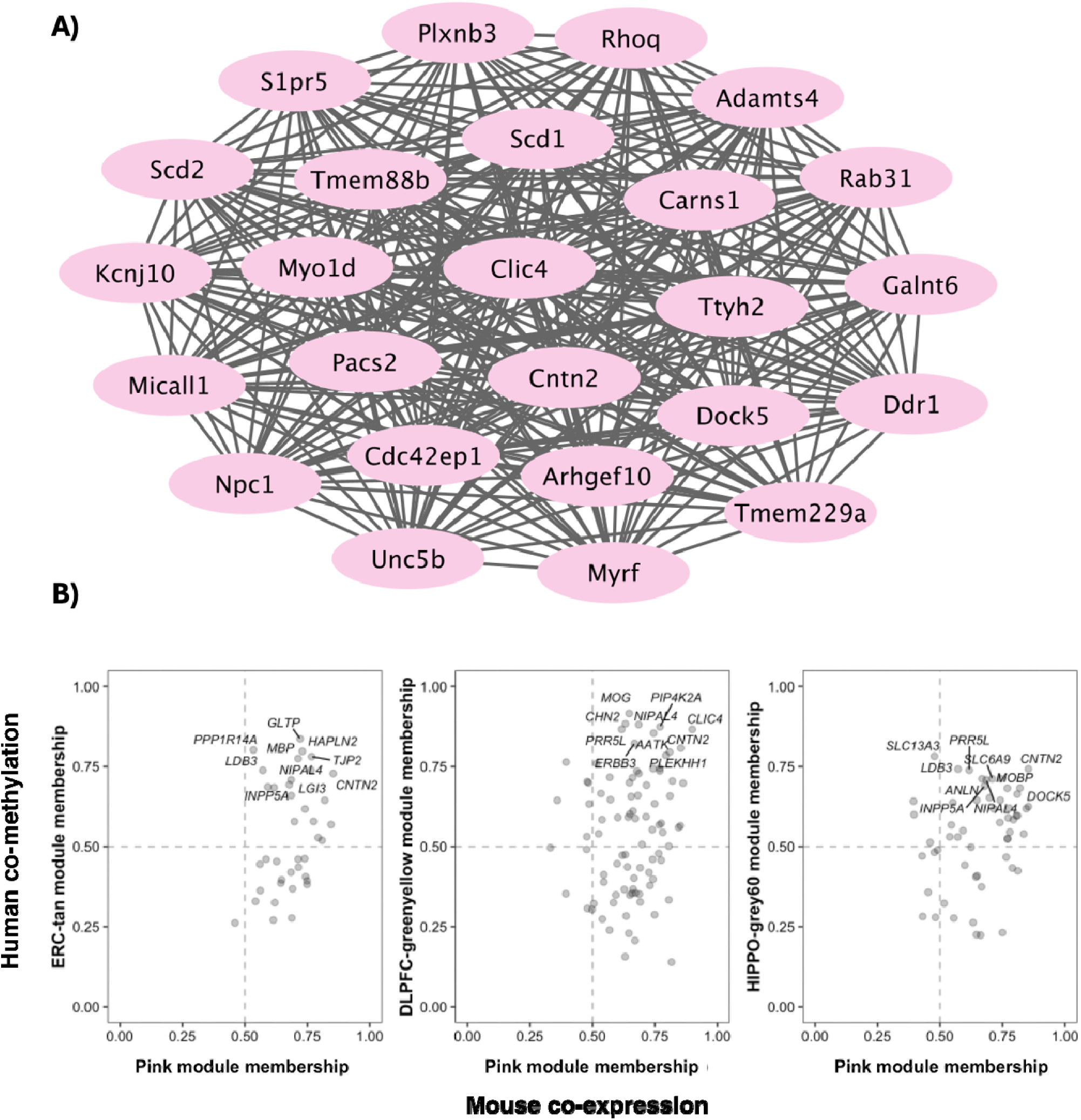
Brain co-expression network analysis of early-stage AD mouse models. A) WGCNA ‘pink’ oligodendrocyte enriched co-expression module. Co-expression network for the ‘pink’ module constructed using Cytoscape software. Nodes represent genes and edge lines represent co-expression connections. The top 25 most connected genes out of the 442 total genes in the module are shown. B) Scatter plots showing the relationship between human co-methylation and mouse co-expression module membership (MM) for genes in the DLPFC-greenyellow, ERC-tan, and HIPPO-grey60 co-methylation modules. The x-axis represents MM in the pink mouse co-expression module; the y-axis represents MM in the human co-methylation modules of interest. Genes with high MM in both species are labelled. Dashed lines indicate the MM threshold of 0.5. DLPFC, dorsolateral prefrontal cortex; ERC, Entorhinal cortex; HIPPO, hippocampus.

**Fig. 6.**
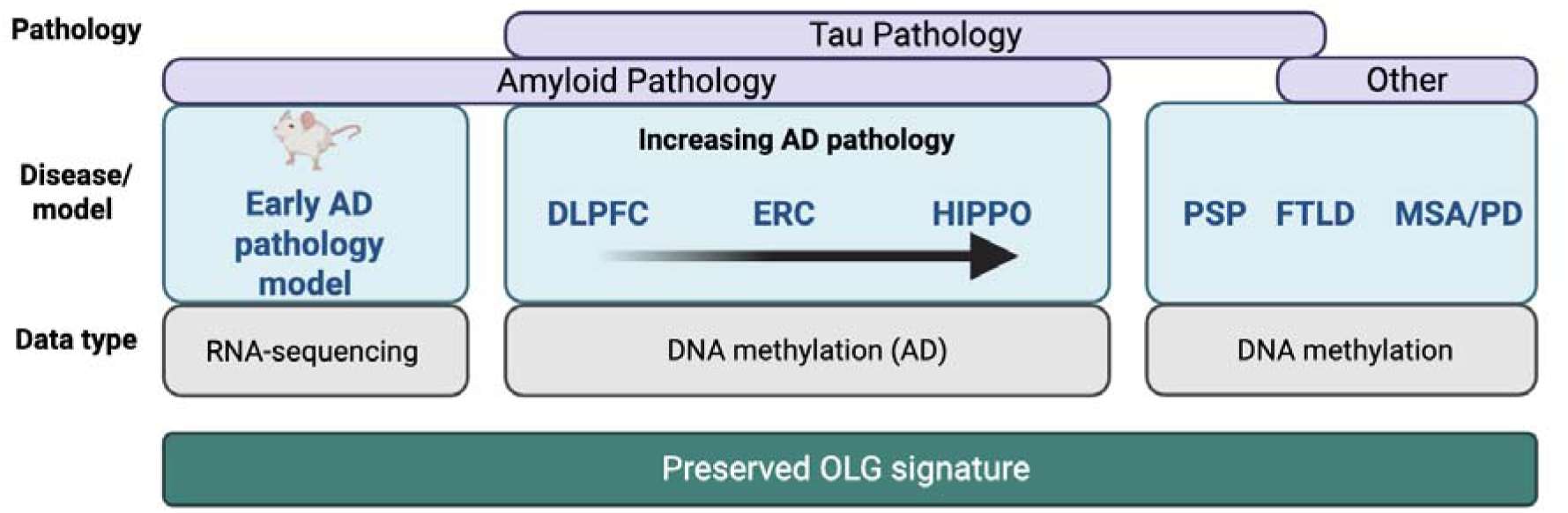
Schematic outlining the datasets and disease contexts in which a preserved oligodendrocyte (OLG) signature across neurodegeneration was identified. AD, Alzheimer’s disease; DLPFC, dorsolateral prefrontal cortex; ERC, entorhinal cortex; FTLD, frontotemporal lobar degeneration; HIPPO, hippocampus; MSA, multiple system atrophy; OLG, oligodendrocyte; PSP, progressive supranuclear palsy; PD, Parkinson’s disease.

## DISCUSSION

We employed network analysis across several DNA methylation and gene expression datasets representing different stages of AD pathology to investigate the effect of DNA methylation on dysregulation of OLG relevant genes in AD. Through these networks, constructed on different modalities of omics data derived from human and mouse brain tissue, we have identified AD-associated OLG signatures shared between more mildly and severely affected human brain regions as well as in mouse models of early stages of AD pathology. Although molecular changes to oligodendrocytes have been reported for neurodegenerative diseases and those with neuropsychiatric symptoms (e.g. PSP[68], schizophrenia[69]), the mechanisms underpinning OLG dysfunction across diseases still needs more investigation. Our findings underpin the importance of studying DNA methylation changes and OLGs as a potentially early contributor to neurodegenerative processes, rather than as passive bystanders, a concept which is in its infancy of being explored[70].

From the human brain co-methylation networks, both the DLPFC-greenyellow and ERC-tan modules had hub genes which are known as important OLG genes and key regulators of myelination: *MOG* and *MYRF* respectively. Strengthening confidence that these modules capture meaningful OLG-associated biology, we found significant positive correlations between DNA methylation and gene expression for these hub genes. *MOG* is a gene expressed by mature OLGs. *MYRF* is a transcription factor that is essential for myelin maintenance[71] and a target of *SOX10*, which is another crucial activator of OLG differentiation-related genes[72]. Recently, using a distinct analysis approach to ours, *MYRF* has been identified as being dysregulated at the DNA methylation and gene expression levels in AD[73]. We noticed several methylation sites with high module membership (MM) across all three co-methylation modules mapped to the gene *ATP11A*, a gene implicated in a hypomyelinating leukodystrophy[63]. Mechanisms behind effects of mutations in this gene were suggested to involve disruption of phospholipids in the cell membrane which led to, amongst other effects, disrupted cell cholesterol homeostasis. Given the known importance of cholesterol functioning in OLGs mediated through the *APOE-*ε4 allele in AD[74], this was of interest. As described, a link between AD and leukodystrophies has been suggested, with several genetic risk factors found to be shared between the two conditions[22]. In a previous transcriptomic study of AD, *ATP11A* was shown to be upregulated in AD compared to controls[75], and we also observed a nominally significant upregulation across multiple OLG clusters in AD compared to controls.

Additional genes with known relevance to AD pathology and white matter perturbations were also present in these dysregulated OLG modules/signatures. The gene *QDPR,* which codes for quinoid dihydropteridine (an enzyme important in the regeneration of tetrahydrobiopterin (BH4), important in nitric oxide production) was found to present across all three modules, and also within disease-associated human snRNAseq and the mouse OLG co-expression modules.

There are some limitations to the use of bulk DNA methylation data in investigating cell-type specific dysregulation, given that cell-type specific methylation changes may be masked by noise from other cell types. However, by using a complementary analysis based on human snRNA sequencing data, we found a significant overlap between genes within the co-methylation OLG modules and those differentially expressed within a cluster of OLGs that had been previously identified as AD associated by Mathys et al[44]. Together with the findings from bulk tissue DNA methylation-gene expression analysis, this supports that DNA methylation acts on the dysregulation of the identified AD-associated OLG signature. We further investigated co-expression modules within the scRNA-seq data and identified that the hd-turquoise module was associated with several pathology measures in the disease associated Oli3 cluster and contained an overrepresentation of genes that we had identified as showing dysregulated DNA methylation. This finding at the single nucleus level reinforces OLG specific observations in the bulk human DNA methylation. To address additional limitations of using bulk human brain tissue (e.g. temporality of events), we further complemented our analyses with co-expression networks in early-stage mouse model data, and identified an amyloid pathology associated module that was enriched for genes showing OLG methylation changes in human AD brain tissue. The finding of cross-species overlap can be interpreted as evidence of shared oligodendrocyte-associated biological processes, though not necessarily direct equivalence of molecular mechanisms. Another limitation of the analysis is the inability to distinguish between 5-methylcytosine (5mC) and 5-hydroxymethylcytosine (5hmC), an oxidative derivative of DNA methylation that is thought to play a role in OLG fate determination[76]. Understanding the role of this specific epigenetic mark could contribute to further insights into the role of DNA methylation on OPC to OLG maturation, which is suggested to be perturbed in AD pathology[77], and warrants further investigation in future studies. AD is frequently accompanied by mixed neurodegenerative and vascular pathologies, some of which may influence OLGs and white matter[78]. As we relied on primary neuropathological classifications provided in the original datasets, detailed stratification by co-pathology was not possible. We therefore acknowledge that mixed pathologies may contribute to some extent to the observed signatures. Likewise, we cannot discard the existent of some degree of AD pathology in the other neurodegenerative diseases used as comparison groups in the preservation analysis. Notwithstanding, the mouse model, which reflects early amyloid-driven pathology in the absence of tau aggregation, complements the human-post mortem data.

## CONCLUSIONS

We have found consistent OLG AD-associated DNA methylation signatures across brain regions affected at different stages of disease progression; the entorhinal cortex and hippocampus are known to exhibit changes early in AD, whilst the dorsolateral prefrontal cortex is typically affected later on in disease[79]. The finding of aberrant DNA methylation across both early- and late stages of pathology suggests alterations contributing to disease early on rather than a secondary effect of neurodegeneration. This interpretation is supported through our complementary analysis of the mouse co-expression modules, which represent an early-stage of amyloid pathology and where the mice have no tau pathology. Furthermore, we saw that there was very high preservation of the same OLG co-methylation AD modules within a primary tauopathy (PSP) dataset and to some extent within datasets consisting of TDP-43 proteinopathies and synucleinopathies as well (**Fig. 6**). Together, these findings highlight oligodendrocyte dysregulation as an important feature of AD, while also possibly suggesting broader relevance across neurodegenerative diseases. Future studies will be essential to determine whether these changes reflect shared mechanisms or disease-specific processes.

In summary, our findings reveal that oligodendrocytes, often overlooked in neurodegeneration, show early and disease-relevant DNA methylation changes and concordant gene expression changes in AD, offering insights into molecular mechanisms of OLG dysfunction that may be common across neurodegenerative disorders.

## Ethics approval and consent to participate

All studies from which datasets were utilised had received local ethical approval and informed consent.

## Consent for publication

Not applicable.

## Availability of data and materials

### Data Availability

Human AD DNA methylation data were accessed through the Gene Expression Omnibus (GEO; accession number **GSE125895**). The independent ROSMAP DNA methylation dataset and RNA sequencing data were accessed through Synapse (ID: syn7357283 and syn3388564, respectively). Mouse RNA-sequencing data are available under GEO accession number **GSE137313**. Human single-nucleus RNA sequencing data were accessed through Synapse (ID: **syn18485175**).

### Code Availability

All code used for data preprocessing, network construction, preservation analysis, and figure generation has been deposited in a publicly accessible GitHub repository: https://github.com/CBettencourtLab/AD_OLGDNAm.

## Author contributions

KF contributed to data analysis, prepared the figures, and drafted the manuscript; HMGS contributed to data analysis, figure preparation and interpretation; UY, IP, MM, JH, TL, RdS and DAS, contributed with datasets and/or interpretation; CB conceptualised and supervised the study; All authors critically reviewed and approved the manuscript.

## Acknowledgements

We would like to thank research groups and initiatives that made datasets used in this research publicly available, as well as all brain donors and their families.

