## Supplementary Fig for "Early oligodendrocyte dysfunction signature in Alzheimer’s disease: Insights from DNA methylomics and transcriptomics"

**Supplementary Figure S1: Human Co-methylation Modules across ERC, DLPFC, HIPPO and CRB**
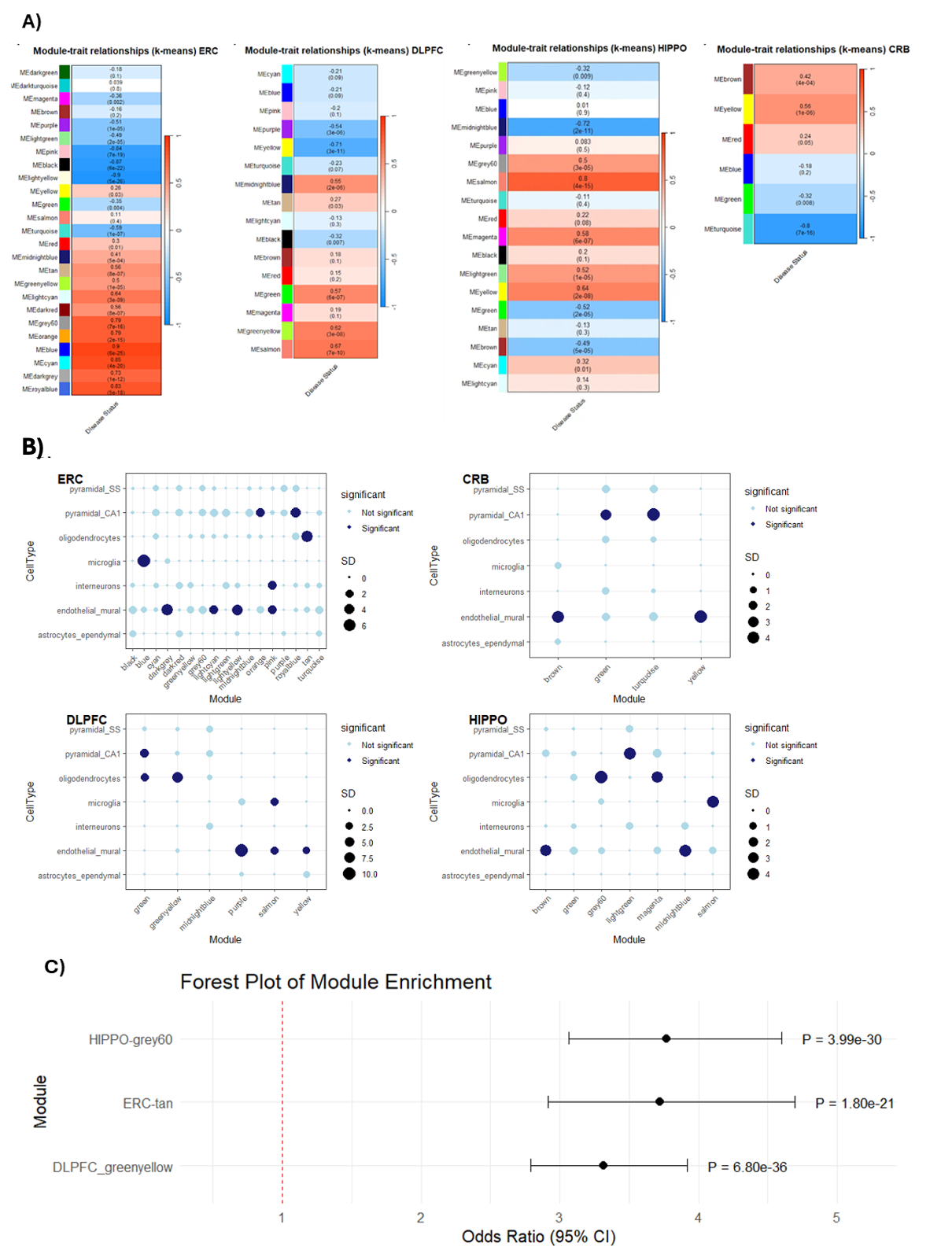

**A) Module trait relationships for ERC, DLPFC, HIPPO, and CRB networks**. The rows represent the co-methylation module eigengenes (ME) and their colours, and the column represents the correlation of the methylation levels of CpGs in each module with the disease status. *p*-values are presented within each cell and the colour scale at the right indicates the strength of the correlation (darker cells depict stronger correlations, with blue representing negative and red representing positive correlations). **B) Enrichment of modules using EWCE for ERC, CRB, DLPFC and HIPPO networks**. Dark filled circles highlight the cell types found to be significantly enriched with adjusted p<0.05 after Bonferroni correction over all cell types within each module; the size of the circles represents the number of standard deviations (SD) from the mean. Cell-type enrichment analysis on the AD-related modules was performed using the package EWCE^52^ and associated single-cell transcriptomic data ^51^.  **C)** **Forest plot of module enrichment for oligodendrocyte (OLG) genes within co-methylation modules using curated OLG gene lists**. Odds ratios (ORs) with 95% confidence intervals are shown for the enrichment of a curated list of OLG genes within AD-associated co-methylation modules derived from three brain regions: DLPFC (greenyellow), ERC (tan), and HIPPO (grey60). EWCE, expression-weighted cell-type enrichment; ERC, entorhinal cortex; DLPFC, dorsolateral prefrontal cortex; HIPPO, hippocampus; CRB, cerebellum; AD, Alzheimer’s disease; OLG, oligodendrocyte.

**Supplementary Figure S2: Human brain AD-associated OLG co-methylation modules**

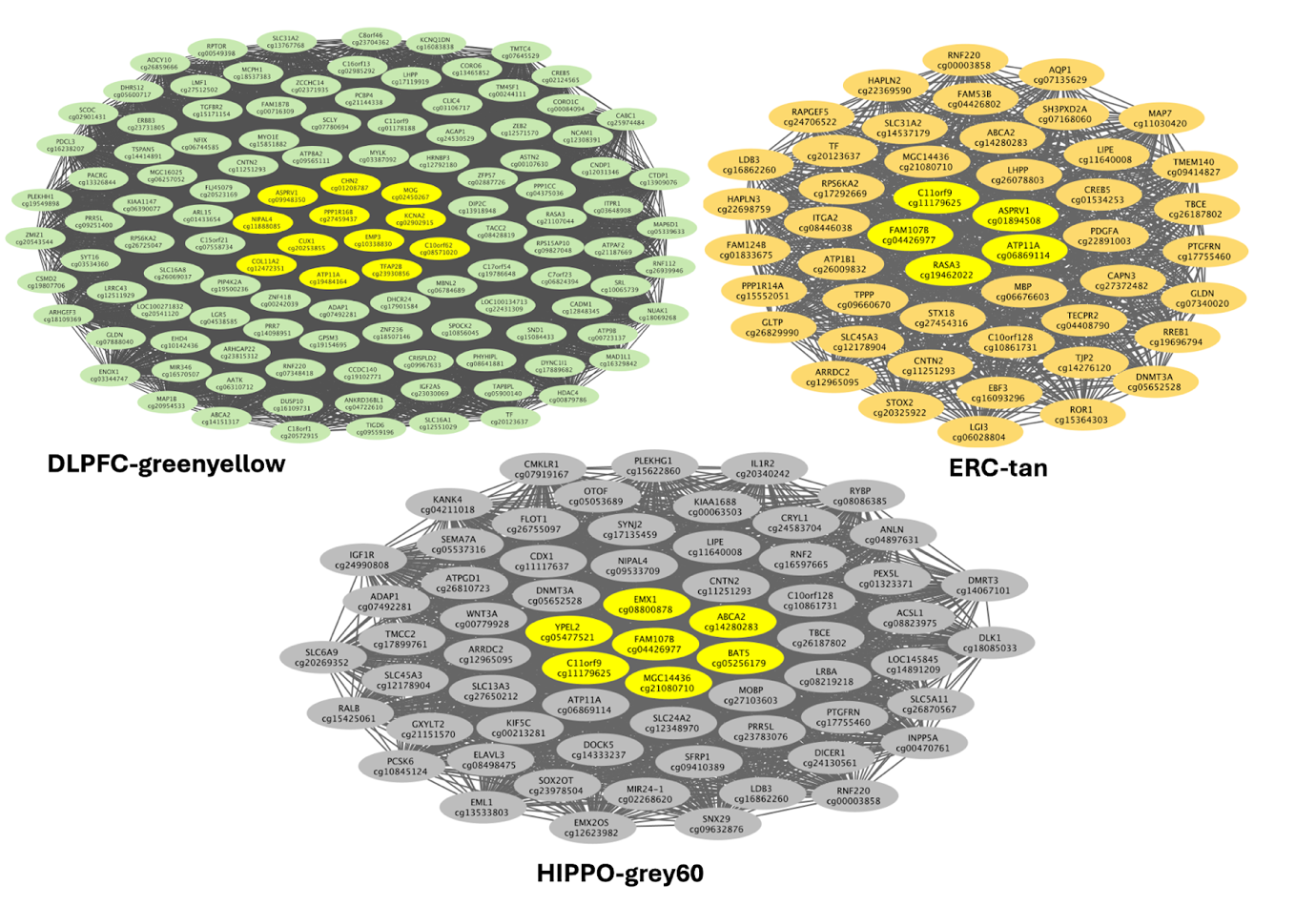

The top 5% MM CpGs and the corresponding genes of the DLPFC-greenyellow, ERC-tan mod and HIPPO-grey60 are shown. Highlighted in yellow as hub genes are the top 0.5% of genes with the highest MM in each module. ERC, entorhinal cortex; DLPFC, dorsolateral prefrontal cortex; HIPPO, hippocampus; AD, Alzheimer’s disease; OLG, oligodendrocyte.

**Supplementary Figure S3: Expression levels of genes present in disease associated co-methylation modules in OLG snRNA-sequencing clusters.**

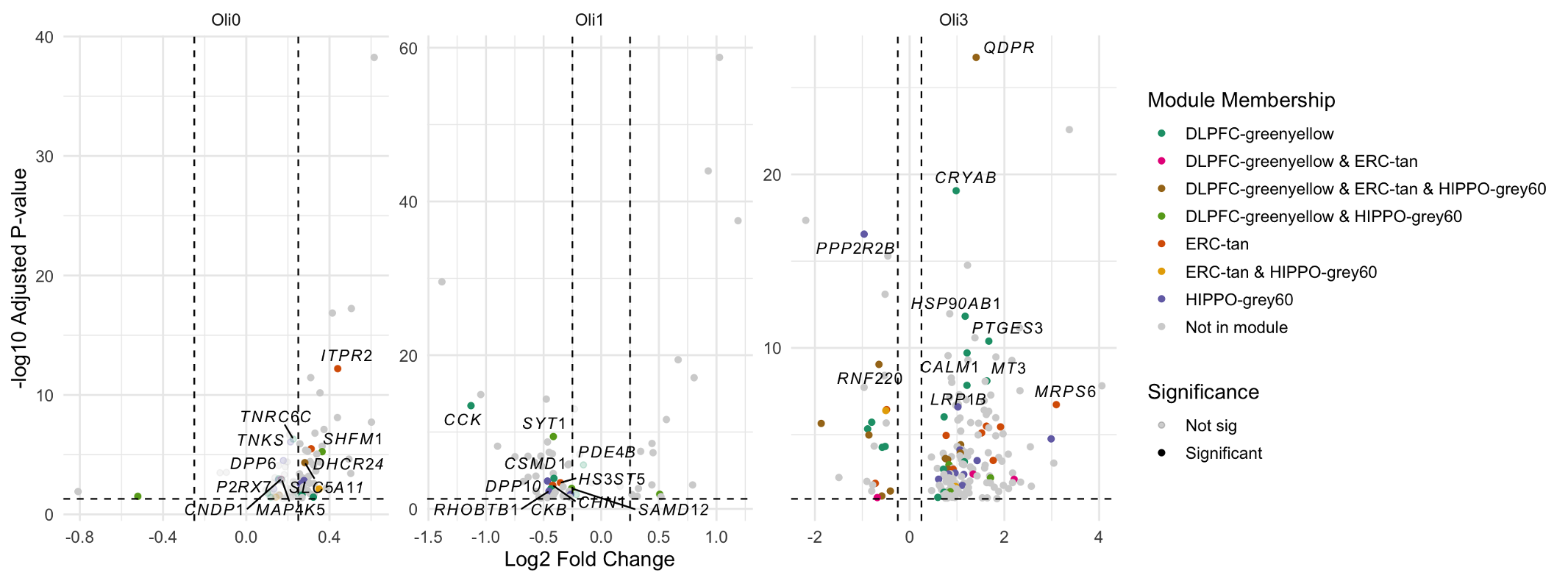

Volcano plots showing differential gene expression across Oli0, Oli1, and Oli3. Significantly differentially expressed genes (adjusted p-value < 0.05) from the co-methylation modules are coloured by their module membership. Genes present in multiple modules are shown in combined colours. Labelled genes are the top 10 most significantly dysregulated within each subcluster which are also module members. AD, Alzheimer’s disease; DLPFC, dorsolateral prefrontal cortex; ERC, entorhinal cortex; HIPPO, hippocampus; OLG, oligodendrocyte.

**Supplementary Figure S4: Mouse brain co-expression modules**

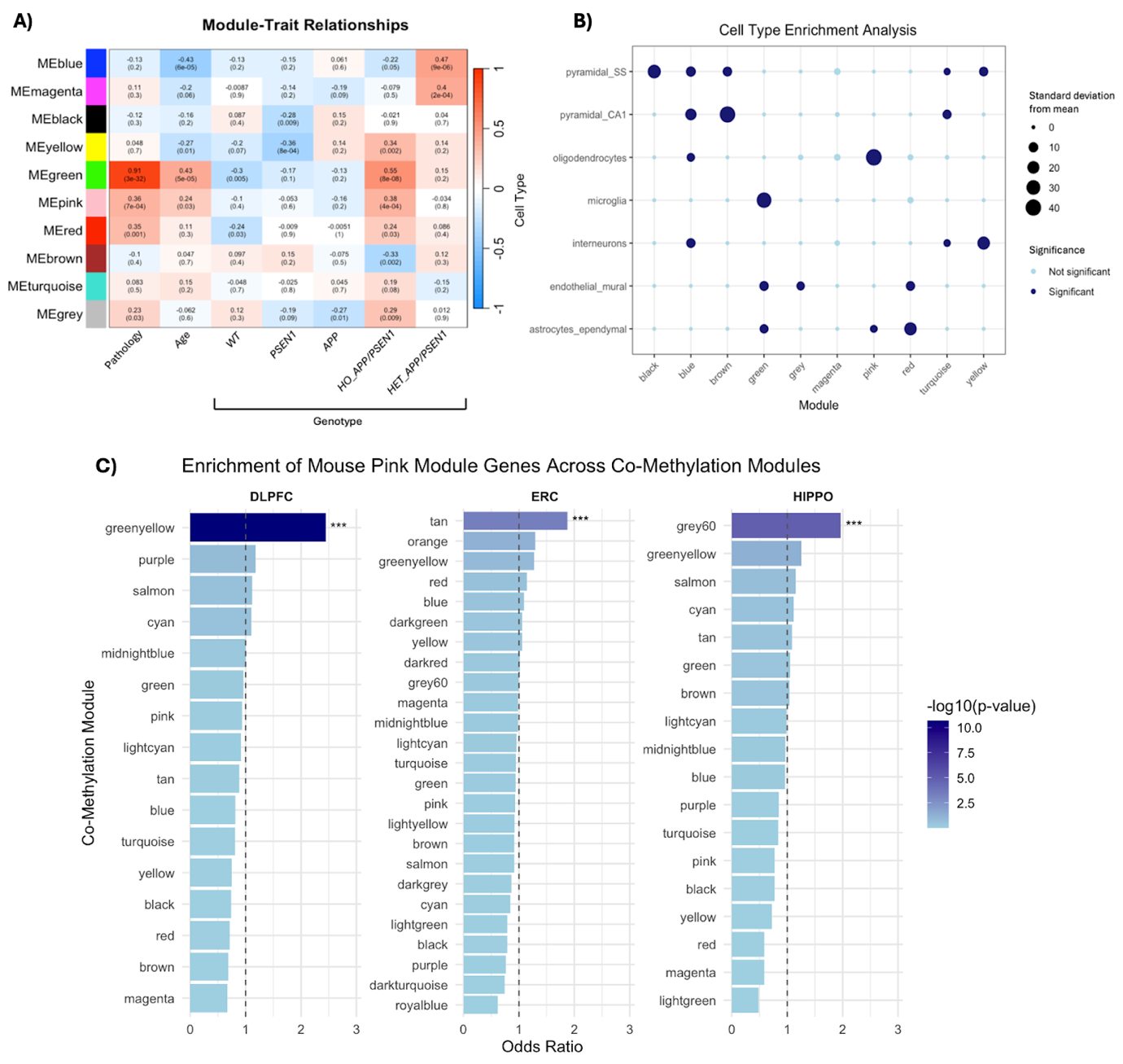

**A) Module trait relationships for mouse AD model co-expression networks.** The rows represent the co-expression module eigengenes (ME) and their colours. The columns represent the correlation of the expression levels of genes in each module with AD related traits: pathology (Aβ presence, as assessed by immunohistochemistry), age, and genotype (*WT, PSEN1, APP,* Heterozygous *APP/PSEN1*, Homozygous *APP/PSEN1*, as described in **Table S1**). The red-blue colour scale represents the strength of the correlation (darker cells depict stronger correlations, with blue representing negative and red representing positive correlations). Following Bonferroni multiple testing correction, the significance threshold was adjusted top ≤ 0.005. P-values are presented within each cell. **B) Enrichment of modules using EWCE for mouse co-expression networks**. The EWCE package^52^ and associated single-cell transcriptomic data^51^ was used to explore whether genes composing each of the 10-co-expression modules identified by WGCNA are relevant for specific brain cell types. Circles filled with dark blue represent significant enrichment of modules with specific cell type signatures using an adjusted p<0.05 threshold after Bonferroni correction over all cell types within each module. Circle size indicates the number of standard deviations from the mean. **C)** Enrichment of mouse pink module genes within human co-methylation modules across brain regions. Bar plots show the odds ratio (x-axis) for enrichment of genes from the mouse pink module within each co-methylation module (y-axis) identified in dorsolateral prefrontal cortex (DLPFC), entorhinal cortex (ERC), and hippocampus (HIPPO) DNA methylation networks. Bars are coloured by the −log₁₀(*p*-value) for enrichment, with darker colours indicating stronger significance. The dashed vertical line at odds ratio = 1 indicates no enrichment. *** denotes *p* < 0.001.
